# Sex moderates the relationship between aortic stiffness, cognition and cerebrovascular reactivity in healthy older adults

**DOI:** 10.1101/2020.02.18.955146

**Authors:** Dalia Sabra, Brittany Intzandt, Laurence Desjardins-Crepeau, Antoine Langeard, Christopher J. Steele, Frédérique Frouin, Richard D. Hoge, Louis Bherer, Claudine J. Gauthier

## Abstract

It is well established that sex differences exist in the manifestation of cardiovascular diseases. Arterial stiffness (AS) has been associated with changes in cerebrovascular reactivity (CVR) and cognitive decline in aging. Specifically, older adults with increased AS show a decline on executive function (EF) tasks. Interestingly, the relationship between AS and CVR is more complex, where some studies show decreased CVR with increased AS, and others demonstrate preserved CVR despite higher AS. We investigated the possible role of sex and hematocrit (HCT) on these hemodynamic relationships. Acquisitions were completed in 48 older adults. Dual-echo pCASL data were collected during a hypercapnia challenge. Aortic PWV data was acquired using cine phase contrast velocity series. A moderation model test revealed that sex moderated the relationship between PWV and CVR and PWV and EF. In addition, there was a significant effect of HCT on the sex differences observed in the moderation effect on EF. Together, our results indicate that the relationships between PWV, CVR and EF are in part mediated by sex and HCT.

**Highlights:**

- This study investigates the role of sex on cerebrovascular measures of brain health and congition
- Sex moderates the relationship between PWV, cerebrovascular reactivity and cognition
- Hematocrit influences the sex differences observed

## 1. Introduction

Cardiovascular diseases (CVD) are the leading cause of death worldwide in males and females, and on average, someone dies of CVD every 38 seconds, resulting in 2,303 deaths per day due to CVDs (Members et al. 2010). Cardiovascular risk is highly sex-dependent, as males exhibit greater incidences and prevalence than females (Petrea et al. 2009; Appelros et al. 2009). However, the risk of heart disease is often underestimated in females due to the misperception that females are protected against CVDs. This could be explained, in part, by the findings of epidemiological studies indicating that premenopausal females are relatively protected from CVDs when compared to age-matched males (Coutinho 2014; Ellekjaer et al. 1997; DuPont et al. 2019). Yet, the incidence of CVD increases disproportionately in females after menopause (Coutinho 2014; Ellekjaer et al. 1997; DuPont et al. 2019), typically 7 to 10 years later, and is the most common cause of death in females over the age of 65 years (Maas et al. 2010). This calls for further understanding of the underlying mechanisms for sex differences involved in the development of cardiovascular diseases. The pathophysiology underlying CVDs is thought to differ depending on the presence of sex hormones leading to differences in vascular properties, including differences in vascular tone (DuPont et al. 2019). Notably, estrogen is thought to have a positive effect on the inner layer of the artery wall, helping to keep blood vessels flexible, and allowing them to relax and expand to accommodate increases in blood flow (Maas et al. 2010; Towfighi et al. 2009). Many of these effects are mediated or influenced by nitric oxide (NO), a highly reactive gaseous mediator that plays a pivotal role in regulating vessel wall homeostasis (Cannon 1998; McNeill et al. 2002). Consequently, a decline in estrogen in postmenopausal females may lead to arterial stiffening (AS) and thus contribute to the increased prevalence of CVD in females in later life (Pepine et al. 2006; Orshal and Khalil 2004; Shaw et al. 2006). Moreover, recent work suggests a potential protective role of testosterone against AS. Specifically, low testosterone levels in males is associated with greater AS (Kyriazis et al. 2011) and augmentation index, an indirect measure of carotid stiffness (Corrigan et al. 2015).

It is well established that elevated AS is an independent predictor of CVD (Mitchell 2009). The elasticity of large arteries allows for the dampening of the arterial pressure waveform, transforming the pulsatile flow at the heart level into steady blood flow into the micro-vessels (Scuteri et al. 2011; Iulita et al. 2018; Badji et al. 2019). Unfortunately, during aging, large arteries (e.g the aorta, the carotids etc) become stiffer and show a reduced capacity to dampen the arterial pressure waveform (Badji et al. 2019). With aging, the elastic properties of blood vessel walls are known to deteriorate (Novak 2012). In particular, the ratio between elastin and collagen changes in favor of collagen, making the vessel stiffer (Novak 2012). Considering the impact of AS on vascular health, non-invasive methods have been developed to measure it, among which pulse wave velocity (PWV) is considered to be the gold standard (Van Bortel et al. 2012).

The impact of AS differs considerably between males and females because of numerous endogenous factors such as the previously mentioned sex hormones and biochemical properties of the arteries (Rossi et al. 2011; Segers et al. 2007). For instance, it has been shown that aged male and female monkeys develop similar levels of AS but a decrease in elastin was noted only in male monkeys (Qiu et al. 2007). In addition, dyslipidemia and glucose contribute to a modest increase in arterial stiffness only in females (Kim et al. 2014). Moreover, it has been shown that the association between AS and mortality is almost two-fold higher in females compared to males after menopause (Regnault et al. 2012), showing probably differences in pathophysiological mechanisms.

However, some of the sex-related differences observed may be in part due to differences in how sex-specific physiological factors (e.g HCT) affect the measures themselves. For example, recent work suggests that hemodynamic measures, such as PWV, may be affected by differences in the concentration of HCT. Males and females typically have different HCT levels (Yip et al. 1984; Vahlquist and Others 1950; Garn et al. 1975), and whole blood viscosity (hematocrit and plasma viscosity) has been shown to be positively correlated with certain cardiovascular disease factors and measures of vascular function (Parkhurst et al. 2012; Bonithon-Kopp et al. 1993; Levenson et al. 1987). Thus, this raises the possibility that a portion of the sex effects observed in the literature could be mediated, in part, by differences in HGB levels.

AS is also associated with downstream organ damage, especially in high-flow organs such as the brain (Pase et al. 2016; Iulita et al. 2018; Mitchell 2009; O’Rourke and Safar 2005; Badji et al. 2019). Indeed, high pulsatile flow following AS may damage cerebral microvessels, thus, leading progressively to changes in cerebral blood flow (CBF) (Singer et al. 2013; Tarumi et al. 2013; Tarumi et al. 2011). Increased stiffness is also associated with changes in cerebrovascular reactivity (CVR), defined as the ability of microvessels to increase blood flow in response to a vasodilatory stimulus (DuBose et al. 2018; Jefferson et al. 2018). It can be hypothesized that increased stiffening of the aorta and higher pulse pressure may lead to damage in downstream vessels from the damaging effects of the pressure amplification caused by large artery stiffness. Therefore, greater artery stiffness may contribute to the impaired ability of the cerebrovasculature to dilate maximally to augment cerebral blood flow (CBF) in older adults. Yet, the literature has found conflicting results, where some have reported reductions in CVR among older adults with greater aortic stiffness using positron-emission tomography (PET) and transcranial doppler (TCD) (DuBose et al. 2018; Jaruchart et al. 2016), while others demonstrate preserved CVR in the presence of higher aortic PWV (Jefferson et al. 2018; Zhu et al. 2013) using arterial spin labeling (ASL), an MRI technique for noninvasive quantification of CBF. Because these imaging modalities (TCD, PET, ASL) are sensitive to CVR arising from different vessel sizes, these results indicate that the relationship between PWV and CVR is complex. With the evidence that sex differences exist in the manifestation of AS, it is unclear if the conflicting results could partly be due to sex-specific characteristics.

Another consequence of a stiffer vascular network is a change in cognitive function (Singer et al. 2014). There is accumulating evidence from cross-sectional studies that AS is associated with the pathogenesis of cognitive decline in both males and females (Singer et al. 2014; Elias et al. 2009; Fukuhara et al. 2006) in age-sensitive domains such as processing speed (PS), verbal memory, and executive functions (EF) (Poels et al. 2007; Watson et al. 2011). Interestingly, associations between PWV, impaired CVR and severity of dementia have also been established (Silvestrini et al. 2006). Indeed, reduced reactivity has been linked to decreased executive functioning, memory, global cognition, and attention outcomes (Haratz et al. 2015). Interestingly, evidence for sex differences has been reported in a number of specific cognitive domains (Halpern and LaMay 2000). Yet, no study to date has investigated the role of sex on the relationship between cognition and neuroimaging markers of brain hemodynamics. Thus, the purpose of this study is to clarify the impact of sex-related differences on the link between PWV, cognitive performance and CVR and the relative contribution of HCT on these relationships.

## 2. Methods

### 2.1 Participants

Fifty-four healthy older adults (17 males, mean age 63 ± 5 years) completed a magnetic resonance imaging (MRI) session. Participants were recruited through a participant database at the Centre de recherche de l’Institut universitaire de gériatrie de Montréal and from Laboratoire D’Etude de la Santé cognitive des Ainés. Inclusion criteria for participation included being in the age range of 55 to 75 years, approval by a geriatrician to participate, non-smoker, no evidence of cognitive impairment as determined through cognitive tests conducted by a neuropsychologist, and MRI compatibility. In addition, the Mini Mental Status Examination (MMSE) was administered, a global cognitive screening tool used for dementia. Participants with scores of less than 26 (out of 30) were excluded (Kurlowicz and Wallace 1999; Gauthier et al. 2015) (no participant was excluded from the sample for this reason). Other exclusion criteria included individuals taking prescription medication known to be vasoactive (e.g. anti-hypertensive drugs, statins, etc.), presence of cardiac disease, hypertension (including use of anti-hypertensive medication), neurological or psychiatric illnesses, diabetes, asthma, thyroid disorders, smoking within the last 5 years, or excessive drinking (more than two drinks per day). All procedures were approved by Comité mixte d’éthique de la recherche du Regroupement Neuroimagerie/Québec and were conducted according to the Declaration of Helsinki. All participants provided written informed consent. From all participants that were recruited, a total of 48 older adults (17 males, mean age 63 ± 5 years) were included in the analysis. The six participants were excluded as outliers since they were more than 2.5 standard deviations above or below the mean of PWV values. Exclusion of these participants did not change the results.

### 2.2 Cognitive Functioning

Cognitive function was assessed with a comprehensive neuropsychological battery consisting of the following cognitive tests: Similarities, Digit Span Backwards, Digit Span forward, Digit Symbol, Color-word interference test (CWIT), and Trail Making Tests, parts A and B (Gauthier et al. 2013; Intzandt et al. 2019).

A composite score for executive function (EF) was calculated using four cognitive tests from the neuropsychological battery that included the CWIT Inhibition and switching conditions, Trail making test part B and the digit span backward. The trail making test part B (TMT-B), the CWIT inhibition condition and the CWIT switching condition were timed in seconds where a low score (faster response) indicates better functioning. The digit span backward was calculated as the number of successful trials where a higher score indicates better EF.

Individual raw scores for each test were transformed into z-scores. The scores that were response time were multiplied by −1 so that a higher EF composite score indicates better cognitive functioning. Cronbach’s alpha was used as a test of reliability to look at the internal consistency for the group of variables. A Cronbach alpha of 0.789 was computed for executive function showing good internal consistency.

### 2.3 Hypercapnia

As previously described (Gauthier et al. 2015; Gauthier et al. 2013; Intzandt et al. 2019), the hypercapnic manipulation was completed with a computer-controlled gas system with a sequential gas delivery circuit (Respiract™, Thornhill Research Inc., Toronto, Canada). The hypercapnic manipulation consisted of two, 2-minute blocks of hypercapnia, with 2 minutes of air before and after each hypercapnia block. End-tidal partial pressure of CO2 (ETCO2) was targeted at 40 mmHg at baseline and 45 mmHg during the hypercapnia blocks. End-tidal partial pressure of O2 (ETO2) was targeted to be 100 mmHg throughout the experiment. Participants breathed through a soft plastic mask that was firmly placed on their face with adhesive tape (Tegaderm 3M Healthcare, St. Paul MN) to ensure that no leaks were present. Participants completed the breathing manipulation once prior to being in the scanner to ensure adequate comfort levels, and once after the MRI session.

### 2.4 MRI acquisition

All acquisitions were completed on a Siemens TIM Trio 3T MRI system (Siemens Medical Solutions, Erlangen, Germany). A 32-channel vendor-supplied head coil was used for all acquisitions. An anatomical 1 mm^3^ MPRAGE acquisition (TR/TE/flip angle = 300 ms/3 ms/90°, 256×240 matrix) was acquired for the registration process from native to standard space, and to measure grey matter partial volume. A fluid attenuation inversion recovery (FLAIR) acquisition with the parameters: TR/TE/flip angle 9000 ms/107 ms/120°, an inversion time of 2500 ms, 512 × 512 matrix for an in-plane resolution of 0.43 × 0.43 mm and 25 slices of 4.8 mm was used to estimate the presence and severity of white-matter hyperintensities. In addition, a pseudo-continuous arterial spin labeling (pCASL) acquisition was acquired, providing simultaneous BOLD contrast using dual-echo readouts (TR/TE1/TE2/flip angle = 2000 ms/10 ms/30 ms/90°) with 4×4×7 mm^3^ voxels, 64 × 64 matrix and 11 slices, post-label delay = 900 ms, tag duration=1.5 s, and a 100 mm gap during a hypercapnia challenge (5 mmHg end-tidal CO2 change, iso-oxic during two, 2 min blocks).

### 2.5 Aortic Exam

As previously described (Gauthier et al. 2015; Intzandt et al. 2019), during the MRI session a thoracic aortic exam was also acquired using simultaneous brachial pressure recording (Model 53,000, Welch Allyn, Skaneateles Falls, NY USA) using a 24-element spine matrix coil. Black blood turbo spin echo sagittal oblique images were acquired to visualize the aortic arch (TR/TE/flip angle: 700 ms/6.5 ms/ 180°, 1.4×1.4 mm^2^ in-plane resolution, 2 slices at 7.0 mm). A perpendicular plane to the ascending and descending aorta was defined from these images. A cine phase-contrast velocity encoded series was collected (TR/TE/flip angle: 28.6 ms/1.99 ms/30°, 1.5×1.5×5.5 mm^3^) during 60 cardiac cycles in three segments, with velocity encoding of 180 cm/s through plane. A series of cine FLASH images were acquired within the same plane with the following parameters: TR/TE/flip angle: 59 ms/3.44 ms/15°, with 1.2 ×1.2 mm^2^ in-plane resolution and a single slice of 6 mm, 60 cardiac phases, acquired in 8 segments.

### 2.6 Data Analysis

Preprocessing of TI-weighted MPRAGE images were done using voxel based morphometry (VBM) in SPM’s Computational Anatomy Toolbox (CAT) 12 (Penny et al. 2011; Ashburner and Friston 2000; Gaser 2016) to segment grey matter, white matter and cerebrospinal fluid (CSF). The registration matrix from T1 space to MNI space was calculated as part of the VBM pipeline Co-registration of native CVR data was done using a non-linear rigid registration with ANTS (Avants et al. 2008) with a b-spline interpolation to bring them from native to individual T1 space. CAT12 was then used to register from T1 to standard space using a Gaussian smoothing kernel of 8 mm and a non-linear registration with 12 degrees of freedom as previously described. (Intzandt et al. 2019)

### 2.7 Resting CBF Analysis

Resting CBF was calculated as previously described (Intzandt et al. 2019). CSF masks were created individually for each older adult to use as a CSF M0 for CBF quantification. 10 voxels were manually chosen in the same axial slice for each participant, within the lateral ventricles. The M0 was then estimated from the control time series and estimated using a monoexponential recovery with a T1 value of 1.65s. Due to varying anatomical structures, each CSF mask was visually inspected to ensure that the region of interest was located in the ventricles. The Bayesian inference for arterial spin labeling MRI toolbox (BASIL) was used for CBF quantification with the following parameters: labeling: cASL/pcASL; bolus duration: constant (1.5s), post label delay: 0.9s; calibration image: average of the control images; reference tissue type: CSF; mask: CSF mask for each participant; CSF TI: 4.3 s; TE:10 ms; T2: 750 ms; blood T2: 150 ms; arterial transit time: 1.3 s, T1: 1.3s, TI blood: 1.65 s, inversion efficiency: 0.85 (Intzandt et al. 2019).

### 2.8 CVR Analysis

CBF-CVR was processed using Neurolens2 (Gauthier et al. 2013; Intzandt et al. 2019). Preprocessing of all raw images included motion correction and spatial smoothing using a 6 mm Gaussian kernel. The CBF signal was isolated from the first series of echoes using linear surround subtraction (Liu and Wong 2005; Gauthier and Hoge 2012; Gauthier et al. 2012; Intzandt et al. 2019). The CBF fractional change during hypercapnia was obtained by fitting a general linear model to the CBF signal and dividing the estimated effect size by the estimated constant term. Glover’s parameters (1999) (Glover 1999) for a single-gamma hemodynamic response function were used when fitting the linear models, which included linear, quadratic, and third order polynomials representing baseline signal and drifts. The CBF percent change obtained was then divided by the average end-tidal CO2 change during the hypercapnia manipulation for each participant to yield CBF-CVR. The baseline CBF was then used to compute absolute values of CBF-CVR.

### 2.9 Vascular lesion quantification

White matter hyperintensity volume (WMH) for the whole brain was quantified semi-automatically. As previously described (Gauthier et al. 2015; Intzandt et al. 2019) visual identification on FLAIR images were completed by a single rater who was blinded to clinical information, which were then delineated using the Jim image analysis package, version 6.0 (Xinapse Systems Ltd, Northants, UK). WMH volume for the whole brain was quantified using tools from the FMRIB Software Library (FSL).

### 2.10 Pulse Wave Velocity Data

The aortic data was analyzed using the ARTFUN software (Herment et al. 2010), where pulse wave velocity in the aortic arch was computed between the ascending and descending aorta from cine phase contrast images. The aortic lumen contours of the ascending and descending aorta were automatically segmented using amplitude images of cine phase contrast series where flow profiles were also estimated. PWV was calculated as described in (Gauthier et al. 2015; Intzandt et al. 2019).

### 2.11 Blood Tests

Before the MRI exam, participants underwent a blood draw. The blood samples were used to test the concentration of hemoglobin and hematocrit (Gauthier et al. 2015).

### 2.12 Statistical Analysis

Statistical analysis of all data was done using IBM SPSS Statistics for Windows, Version 24.0 (IBM Corp., Armonk, NY). Descriptive statistics for age, education, MMSE scores, WMH, CBF-CVR, PWV and executive functioning scores are reported in the whole sample and compared between males and females in *Table 1*. Statistical comparisons between males and females were done using independent samples *t*-tests. Moderation analyses were performed using the PROCESS Macro for moderation analyses (Hayes 2017). The analyses were bootstrapped to amend any shortcoming in power by simulating greater data based on an algorithm to maintain the current pattern. By default, bootstrapped samples were set to simulate 5,000 samples (Hayes 2017). Moderation effects of sex on the PWV-CVR, PWV-EF, and CVR-EF relations were tested controlling for age and WMH volume. In addition, a moderated moderation analysis was conducted to see if the sex effects in the moderations could be explained by differences in hemoglobin (Fig. 7, Fig. 8).

**Table 1:** Participant Demographics.

| Demographic | All (n=48) | Males (n=17) | Females (n=31) |
| --- | --- | --- | --- |
| Age (years) | 63.35 ± 4.86 | 64 ± 4.37 | 63 ± 5.14 |
| Education (years) | 16.29 ± 3.56 | 16.18 ± 3.32 | 16.3 ± 53.74 |
| EF*<br>(composite score) | 0.003 ± 0.74 | -0.31 ± 0.87 * | 0.177 ± 0.60 * |
| PWV (m/s) | 8.70 ± 2.89 | 8.89 ± 2.99 | 8.59 ± 2.88 |
| MMSE (out of 30) | 28.79 ± 0.94 | 28.53 ± 1.32 | 28.94 ± 0.63 |
| Log WMH volume | 0.35 ± 0.15 | 0.36 ± 0.13 | 0.35 ± 0.17 |
| Resting CBF (ml/100g/min)* | 42.46 ± 10.05 | 35.74 ± 8.71* | 46.14 ± 8.85* |
| CBF-CVR<br>(ml/100g/min/mmHg CO<sub>2</sub> ) | 4.64 ± 2.39 | 4.50 ± 2.52 | 4.71 ± 2.36 |
| HBG (g/L)* | 139.49 ± 11.27 | 148.41 ± 11.24 * | 134.45 ± 7.77 * |
| HCT * | 0.421 ± 0.03 | 0.447 ± 0.321* | 0.406 ± 0.241* |

\* $p < 0.05$

### 2.13 Data and code availability

The preprocessing scripts along with scripts used to analyze data can be made available upon reasonable request. Research data is available upon request.

## 3. Results

A total of 48 older adults (31 females, 17 males) were included in the analysis. Participant characteristics are summarized in Table 1. It was found that females had a significantly higher resting CBF (p < 0.05) in whole brain grey matter, higher composite scores for executive functioning (p < 0.05) and lower hemoglobin and hematocrit than males (p < 0.05). There was no difference between males and females for PWV (p > 0.05) and CBF-CVR (p > 0.05) in whole brain grey matter.

### 3.1 Moderation Analysis

Our results revealed a significant standardized direct effect of PWV on CVR (β = 1.6307, SE = 0.4839, 95% CI [0.654, 2.607], p = 0.0016) as depicted in *Figure 1*. The moderation effect (SEX *PWV) was also a significant predictor of CVR (β = −1.013, SE = 0.2957, 95% CI [-1.610, −0.4169], p = 0.0014) showing that the effect of PWV on CVR was a function of sex. Further analysis revealed that the effect of PWV on CVR was significantly positive in males (β = 0.6170, SE = 0.2184, 95% CI [0.1762, 1.0577], p = 0.0072) and significantly negative in females (β = −0.3967, SE = 0.1902, 95% CI [-0.7805, −0.0129], p = 0.0431) (*see Figure 2*).

**Figure 1:**
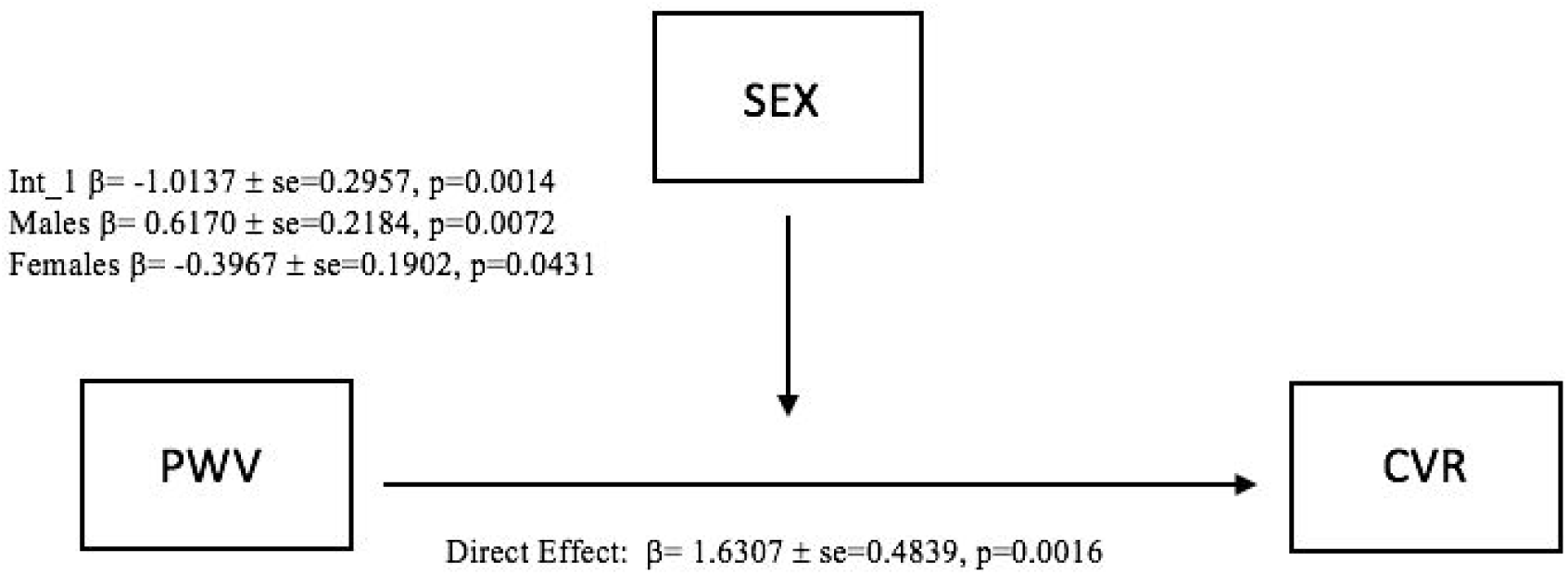
Moderation Effect (PWV*SEX) on CVR.

**Figure 2:**
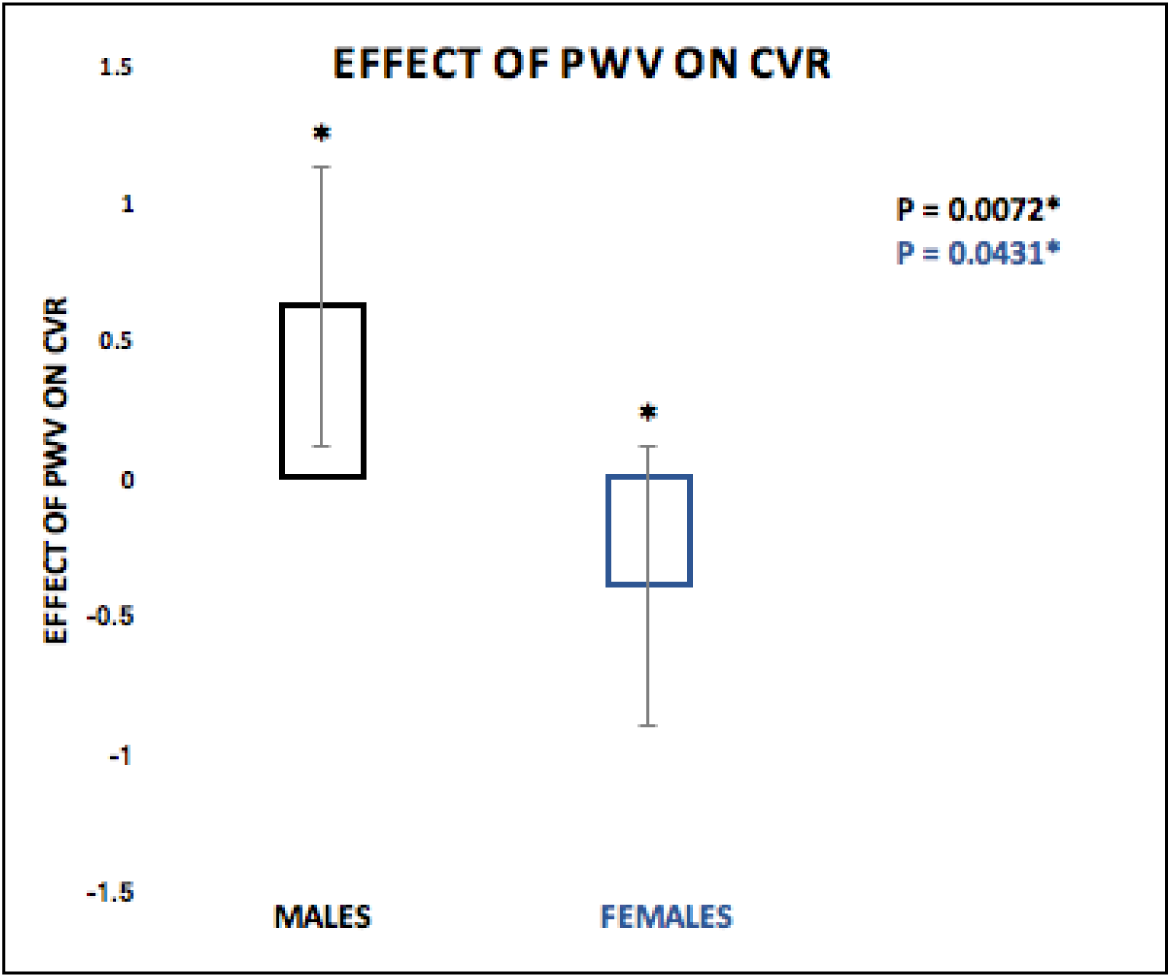
Effect of PWV on CVR. CVR was significantly positive in males (p=0.0072)* and significantly negative in females (p=0.0431)*.

Our results also revealed a significant standardized direct effect of PWV on EF (β = −0.9980, SE = 0.3463, 95% CI [-1.6970, −0.2990], p = 0.0062). The moderation effect (SEX *PWV) was also a significant predictor of EF (β = 0.4479, SE = 0.2117, 95% CI [0.0207, 0.8751], p = 0.0403) showing that the effect of PWV on EF was a function of sex (*Figure 3*). As shown in *Figure 4* the effect of PWV on EF was significantly negative in males (β = −0.5501, SE = 0.1563, 95% CI [-0.8656, −0.2346], p = 0.0011) but not significant in females (β = −0.1022, SE = 0.1361, 95% CI [-0.3769, 0.1725], p = 0.4569).

**Figure 3:**
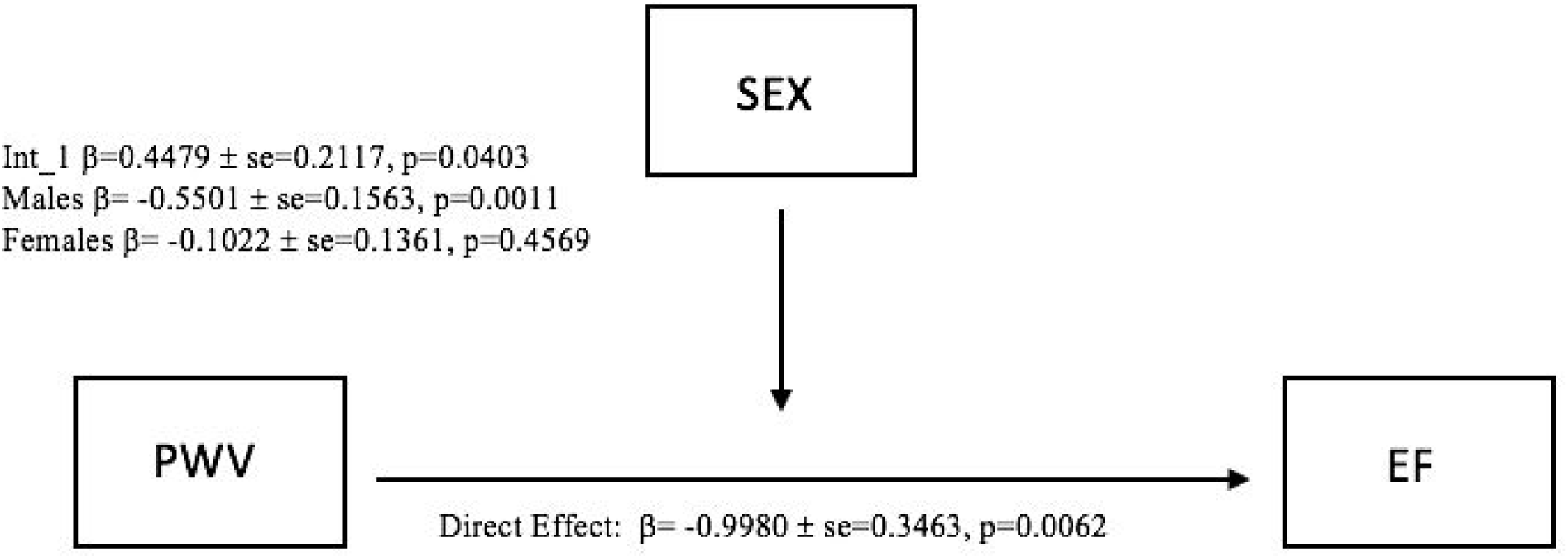
Moderation Effect (PWV*SEX) on EF.

**Figure 4:**
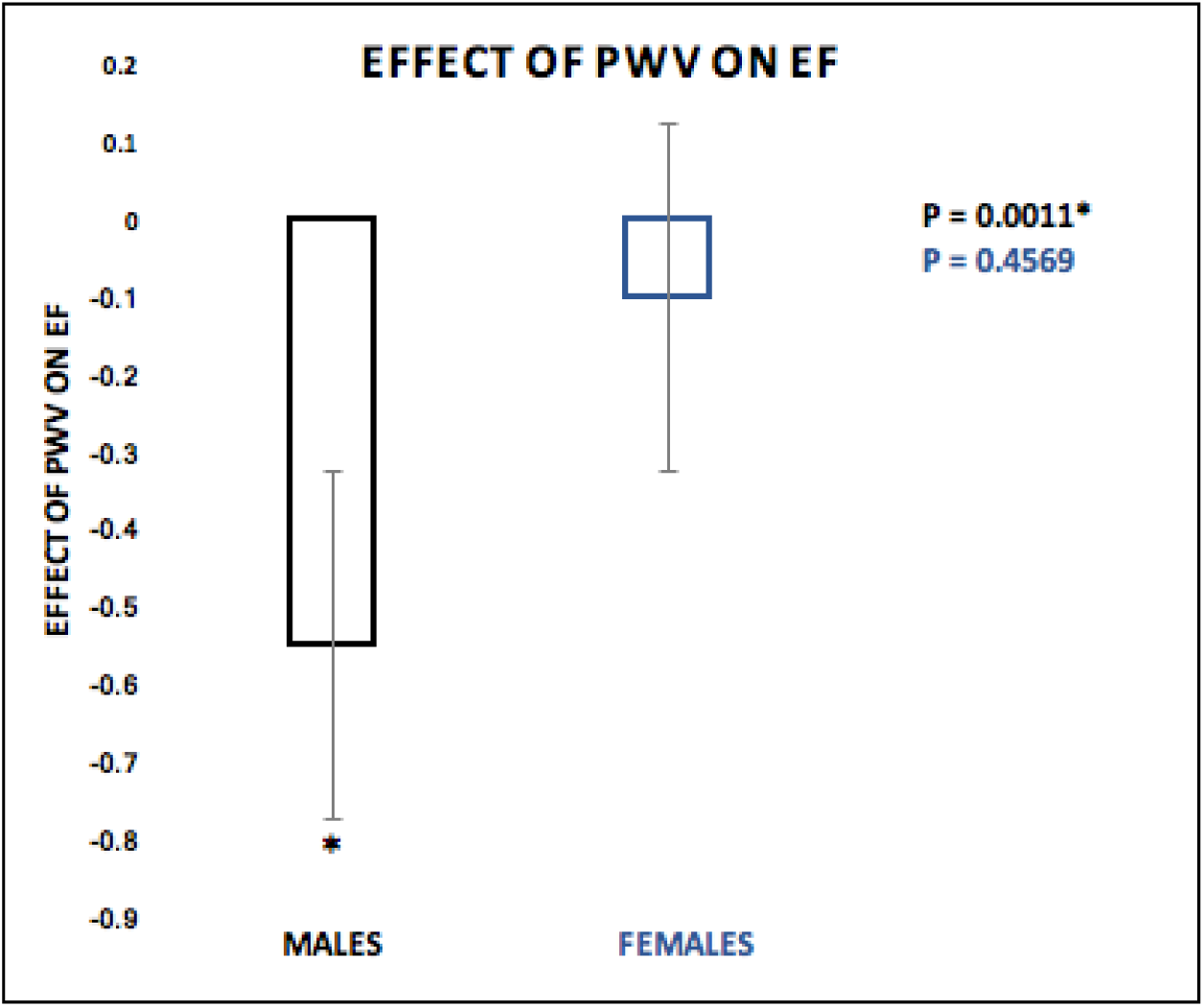
Effect of PWV on EF. The effect of PWV on EF was significant negative in males (β=-0.5501, SE = 0.1563, 95% CI [-0.8656, −0.2346], p=0.0011) but not significant in females (β = −0.1022, SE = 0.1361, 95% CI [-0.3769, 0.1725], p = 0.4569).

Finally, a moderation analysis also revealed a significant standardized direct effect of CVR on EF (β = −0.8472, SE = 0.3332, 95% CI [-1.5195, −0.1748], p = 0.0148). However, the moderation effect (SEX *CVR) did not predict EF (β = 0.3438, SE = 0.1990, 95% CI [-0.0579, 0.7455], p = 0.0914) *(Figure 5)*.

**Figure 5:**
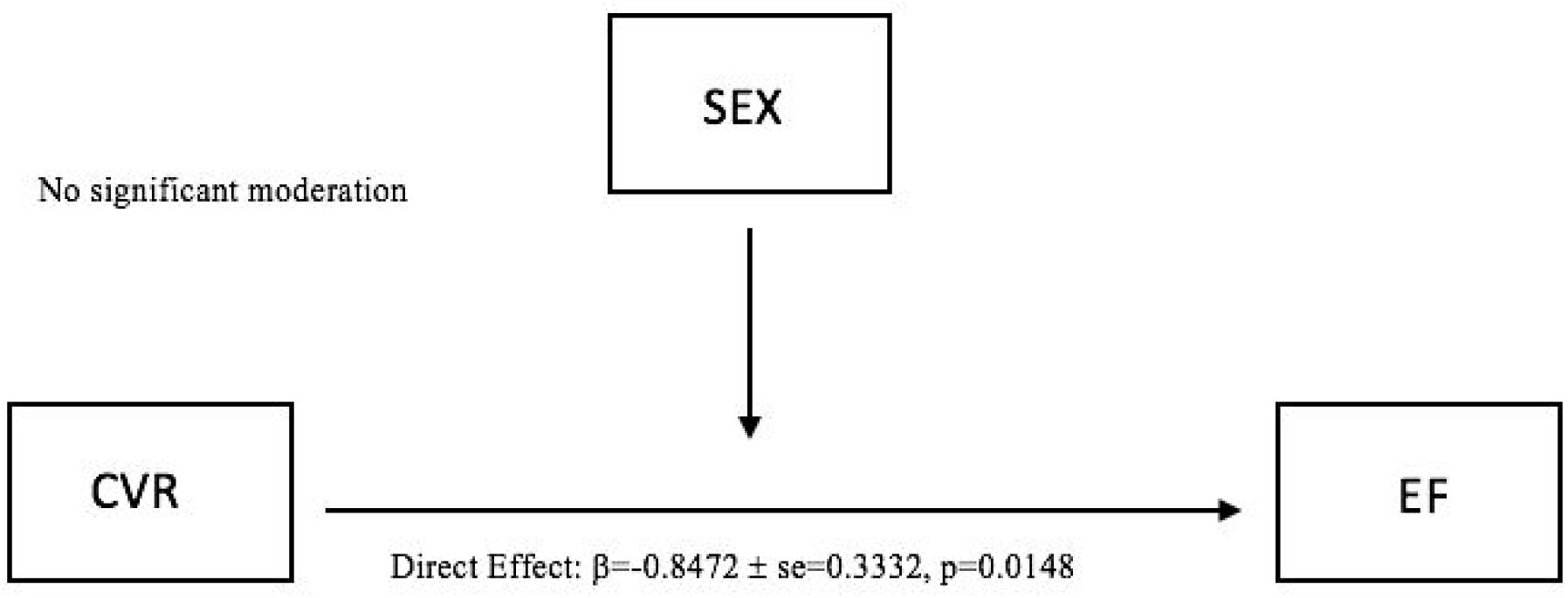
Moderation model (CVR*SEX) on EF.

### 3.2 Moderated Moderation Analysis

A moderated moderation analysis was conducted to determine if the sex effects in our moderation could be explained by differences in, given the sex differences in hematocrit concentration shown in *Table 1.*We found that there was no effect of HCT on the sex differences observed in the moderation effect (PWV*SEX) on CVR (*Figure 6*).

**Figure 6:**
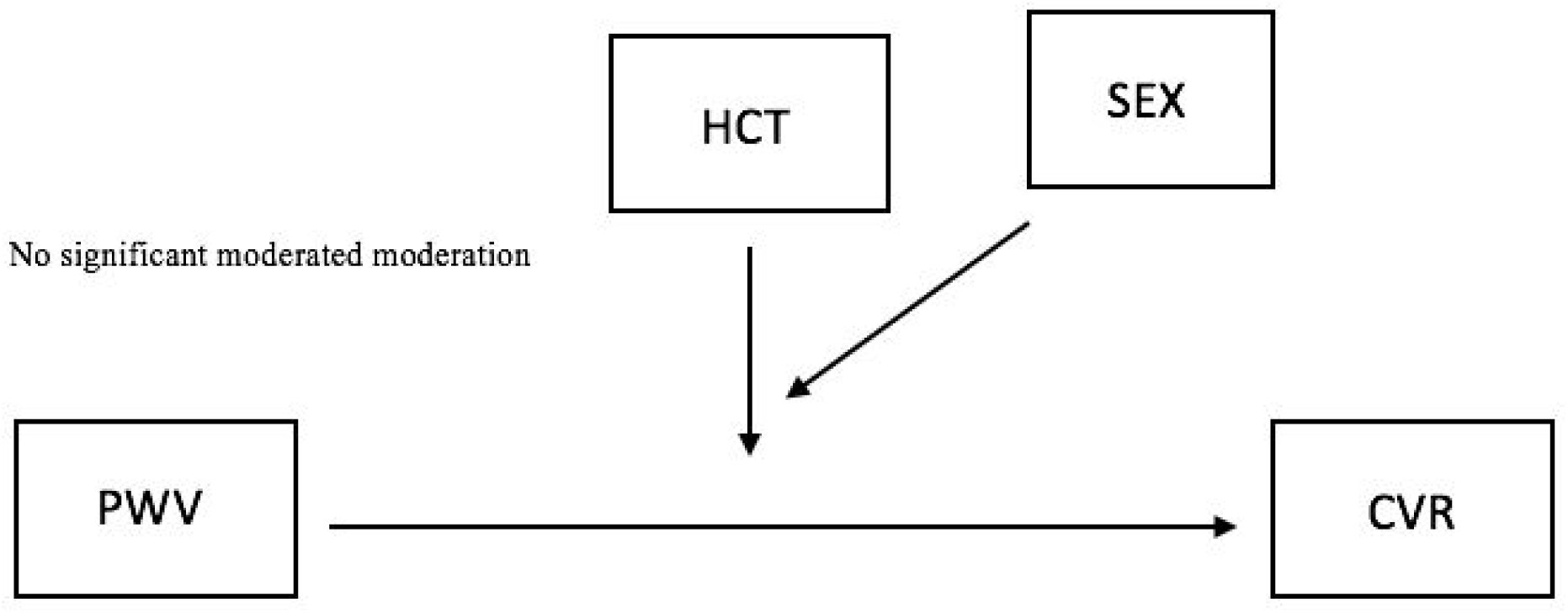
Moderated moderation model depicting the effect of HCT on the relationship PWV on CVR among sexes.

**Figure 7:**
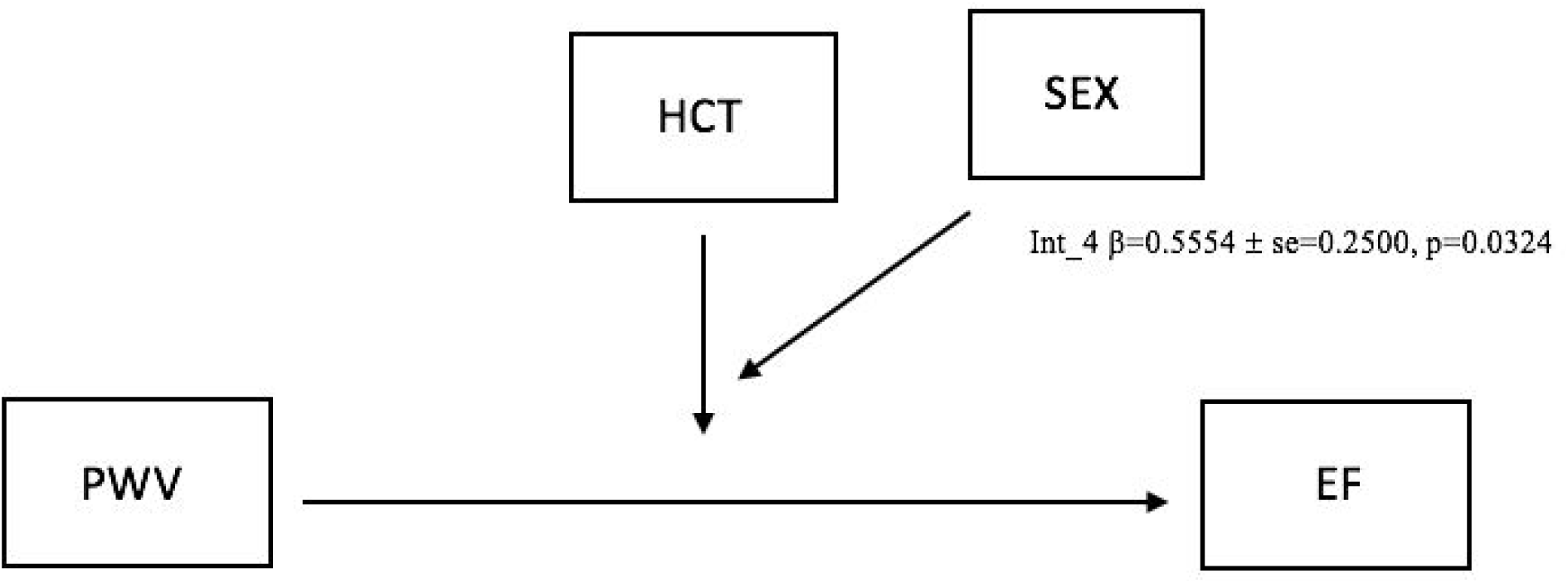
Moderated moderation model depicting the effect of HCT on the relationship PWV on EF among sexes.

**Figure 8:**
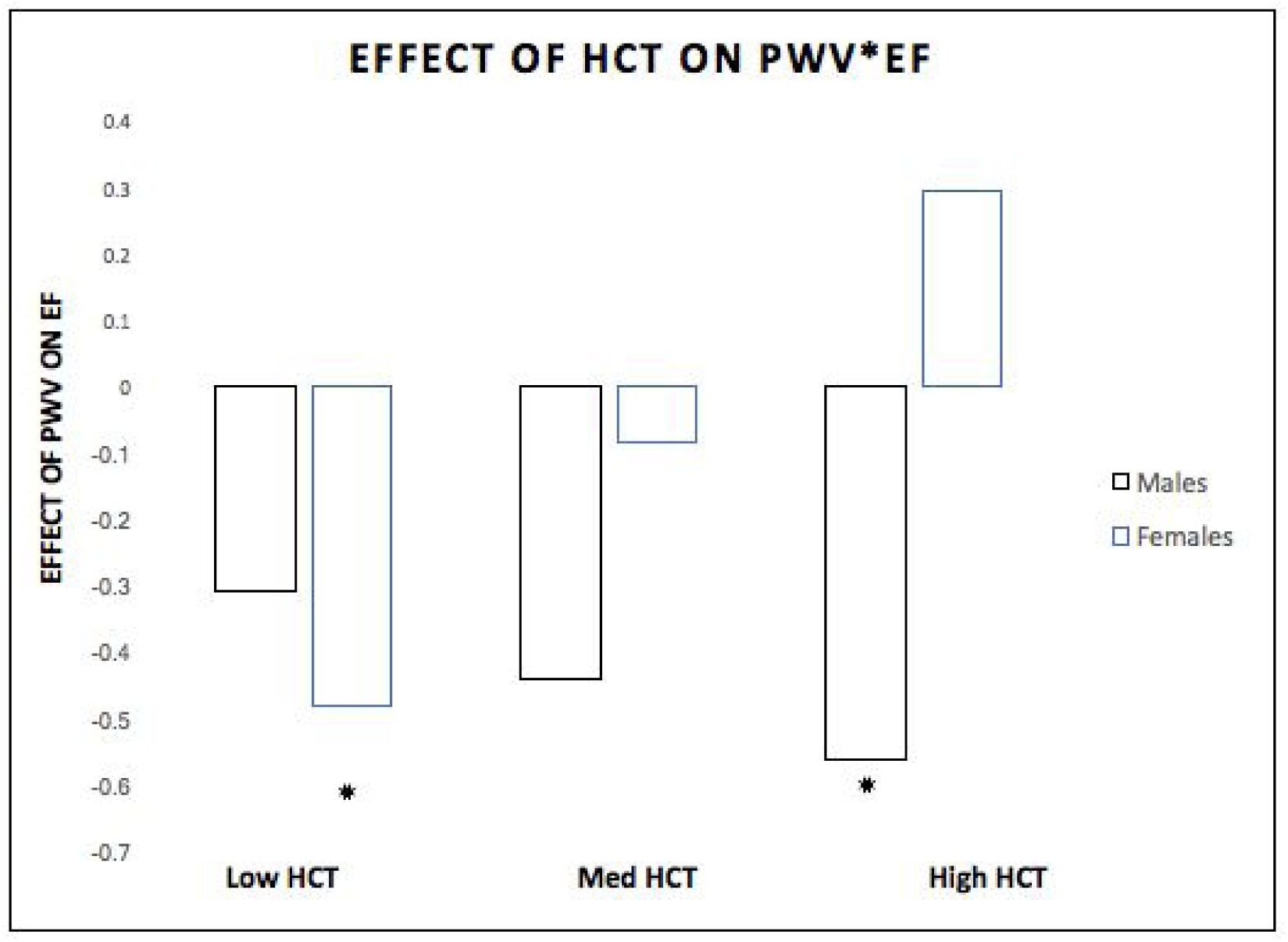
The effect of HCT on the relationship PWV on EF among sexes. The effect of PWV on EF was significantly negative (*p*=0.0206)* in females (blue) with low HCT levels and significantly negative (*p*=0.0030)* in males (black) with high HCT levels.

In addition, through a moderated moderation analysis we tested whether sex moderated the moderation effect of HCT on the relationship between PWV and EF. It was found that the moderated moderation effect (PWV*HCT*SEX) was a significant predictor of EF (β=0.5554, SE = 0.2500, 95% CI [.0492, 1.0615], p=0.0324) (*Figure 7).*As shown in *Figure 8* the effect of PWV on EF was negative in males (β=-0.3097, SE = 0.4479, 95% CI [-1.2164, 0.5970], p=0.4934) and significantly negative in females (β=-0.4797, SE = 0.1985, 95% CI [-0.8815, −0.0779], p=0.0206) with low HCT levels (0.3884 g/L). In addition, the effect of PWV on EF was negative for both males (β=-0.4394, SE = 0.2936, 95% CI [-1.0338, 0.1549], p=0.1427) and females (β=-0.0834, SE = 0.1273, 95% CI [-0.3411, 0.1742], p=0.5161) with medium HCT levels (0.4200 g/L) but did not reach significance. Lastly, it was found that the effect of PWV on EF was significantly negative for males (β=-0.5626, SE = 0.1774, 95% CI [-0.9216, −0.2035], p=0.0030) and positive for females (β=0.2928, SE = 0.2042, 95% CI [-0.1207, 0.7063], p=0.1599) with high HCT levels (0.4500 g/L).

## 4. Discussion

### 4.1 Main Results

In this study, we investigated the impact of sex on the link between PWV, cognitive performance and CVR. An important finding is that sex moderates the relationship between: i) PWV and CVR; and ii) PWV and EF; but not between iii) CVR and EF. Specifically, the effect of PWV on CVR was significantly positive in males and significantly negative in females. Additionally, the effect of PWV on EF was significantly negative in males but not significant in females. Furthermore, results from the moderated moderation analysis revealed that there was no effect of HCT on the sex differences observed in the moderation effect of sex on the relationship between PWV and CVR, however there was a significant effect of HCT on the sex differences observed in the moderation effect of sex on the relationship between PWV and EF. Together, our results indicate that some of the complex relationships between PWV, CVR and EF shown in the literature are being driven by sex and that HCT may be involved in driving some of these sex effects.

### 4.2 Sex Differences

The sex differences identified in this study are consistent with the existing literature, including differences in CBF, HCT and EF performance. In this sample, females have higher resting CBF than males. These findings are similar to previous studies with healthy older adults indicating that females typically display greater resting global CBF (Rodriguez et al. 1988; Esposito et al. 1996) and higher CBF velocities compared to males (Vriens et al. 1989; Martin et al. 1994; Oláh et al. 2000; Tegeler et al. 2013). In addition, several studies have reported data suggesting that males and females tend to present with different levels of performance in certain cognitive domains (Burstein et al. 1980; Kennison 2003; Castonguay et al. 2015). Indeed, our findings of decreased executive function scores in older males compared to older females are also in line with previous research (Halpern and LaMay 2000). Finally, in our population, females had significantly lower hemoglobin levels than males, consistent with previous findings showing that females have mean levels of approximately 12% lower than males (Murphy 2014).

### 4.3 Arterial stiffness and cerebrovascular reactivity

Overall, there is substantial evidence supporting an effect of sex hormone on vascular stiffness (Ogola et al. 2018; DuPont et al. 2019). Over the lifespan, arterial stiffness increases linearly in both males and females however there is a more rapid increase in stiffness in females due to depletions in estrogen levels post menopause (Mitchell 2014; DuPont et al. 2019). Indeed, several studies have shown that hormone receptors, including estrogen and testosterone are cardio-protective (DuPont et al. 2019; Wu et al. 2014; Karas et al. 1994; Dockery et al. 2003). Previous work highlighting the protective effects of estrogen have shown that arterial stiffness, measured using cfPWV, is reduced in postmenopausal females taking hormone replacement therapy (HRT) when compared to matched females not taking HRT (Rajkumar et al. 1997; DuPont et al. 2019). In addition, testosterone was once perceived to play a role in promoting CVD among males (Thompson et al. 1989; Sullivan et al. 1998), however recent epidemiological studies point to the contrary. Testosterone deficiency has been associated with increased cfPWV among healthy older males compared to age-matched males (Vlachopoulos et al. 2014; DuPont et al. 2019). In addition, low testosterone has also been associated with impaired microvascular function and arterial elasticity, measured using the augmentation index. (Corrigan et al. 2015; DuPont et al. 2019). Finally, it is known that testosterone can be converted to estrogen in the brain, so that the protective effects of estrogen may in fact be more preserved in older males than females (Robison et al. 2019). Overall, ample data suggest that sex hormones substantially impact the manifestation of arterial stiffness among males and females. As such, a thorough understanding of the underlying mechanisms that contribute to these sex differences could aid in developing sex-specific strategies to reduce or prevent CVD risk.

Although quantitative measurements of CBF and CVR in relation with arterial stiffness have been performed by others (DuBose et al. 2018; Jefferson et al. 2018), the literature shows conflicting results. While more recent reports suggest a preserved CVR with increased aortic stiffness (Jefferson et al. 2018; Zhu et al. 2013), earlier studies using PET and TCD rather than MRI did not find such an association, but rather reductions in CVR among adults with greater aortic stiffness (DuBose et al. 2018; Jaruchart et al. 2016). It is possible however that these counterintuitive findings could stem from differences in study designs, heterogeneity of target populations (males vs females) and/or differences in imaging modalities. For example, one previous TCD study describing sex-related differences in cerebral vasomotor reactivity has shown increased vasodilatory response in females compared with male subjects (Matteis et al. 1998). More importantly, these studies did not control for menstrual phase or account for sex hormones, which are known to acutely alter CBF and cerebrovascular responsiveness (Brackley et al. 1999; Krejza et al. 2001; Krejza et al. 2003; Krejza et al. 2013; Nevo et al. 2007). Following these and our results, it is imperative that different models should be created for sex rather than just regressing out the effects of sex.

Our work presents the first evidence of a sex moderation in the relationship between PWV and CVR. Our findings of low CVR among females and high CVR among males with increased arterial stiffness are in line with the contradictory results found in the extant literature, with females following the trend observed by DuBose et al and Jaruchart et al. (DuBose et al. 2018; Jaruchart et al. 2016) and male data being in agreement with the results of Jefferson et al. (Jefferson et al. 2018). The biological underpinnings of these differences may be of several origins and as yet there is little data to support a particular hypothesis. However, it could be differential remodeling or damage as a function of stiffness may depend on the time course of stiffness (with men having a potentially more protracted time course since they do not undergo menopause), or may be due to different biases in the CVR technique in males and females. ASL is highly dependent on transit time of the tag for example, and differences in velocity and transit time could for example explain some of these effects. More studies investigating potential sex-dependent methodological biases, and the biological changes that underpin vascular changes in aging in both sexes are needed to start addressing this question.

### 4.4 The association between pulse wave velocity and cognitive function relative to sex

In the present study the relationship between arterial stiffness and EF was in part driven by differences in sex. We also demonstrate that higher arterial stiffness, as measured by PWV, is associated with poorer performance on EF tasks among males only. Our findings are in line with prior studies that have demonstrated sex differences in the relationship between arterial stiffness and cognitive function. For instance, Singer et al. found no significant association between arterial stiffness and global cognition, however negative associations between PWV and composite global cognition were observed in males only when stratifying the sample by sex (Singer et al. 2013). Similarly, Waldstein et al reported males as scoring significantly lower than females on tests of verbal memory at higher levels of pulse pressure, an alternative measure of arterial stiffness (Waldstein et al. 2008). In addition, there is good agreement in the literature showing that arterial stiffness and related central hemodynamics are associated with reductions in cognitive performance on memory, processing speed, and executive function tasks (Iulita et al. 2018; Singer et al. 2014; Badji et al. 2019). However, these studies are limited because they have used a nonspecific measure of cognition (i.e MMSE). Here, we extend their findings to include executive function domains that are especially susceptible to cardiovascular disease (Gorelick et al. 2012). Indeed, our results are consistent with other studies pointing to a general association between PWV and executive dysfunction (Lim et al. 2016; Poels et al. 2007; Hajjar et al. 2016). For example, Hajjar et al found that subjects with higher PWV had the greatest 4-year risk of decline in executive function (Hajjar et al. 2016). It has been hypothesized that this may be due to the fact that the dorsolateral prefrontal cortex, essential for executive function tasks, is situated in a watershed region of the brain (Suchy 2015; de la Torre 2002). Thus, one could speculate that with increased arterial stiffness, these regions may be deprived of perfusion and therefore oxygen earlier than better perfused regions (Suchy 2015; de la Torre 2002).

Our results showed no significant association between arterial stiffness and EF among females. Moreover, females in this cohort displayed significantly better performance than males on the cognitive tasks assessed. It is well known that there are significant differences between sexes in regards to vascular function and cognitive performance among older adults (Castonguay et al. 2015; Narkiewicz et al. 2006). Indeed, across the lifespan, males have a higher risk for CVDs and females are relatively protected until menopause after which they catch up to males, likely due to estrogen depletion (Singer et al. 2013; Narkiewicz et al. 2006). In addition, males have a higher prevalence of mild cognitive impairment compared to females (Petersen et al. 2010; Brodaty et al. 2013; Singer et al. 2013). A possible hypothesis for these sex differences is that males experience cognitive decline earlier in life, while females transition from normal cognition to impaired cognition in the decades following menopause (Petersen et al. 2010).

### 4.5 The association between cognitive function and cerebrovascular reactivity relative to sex

While the vessel reactivity-cognition link has been investigated in cardiovascular and neurodegenerative disorders, the association between CVR and cognition in healthy aging is unclear. Most reports indicate that blood vessel reactivity in the brain decreases with healthy aging (Reich and Rusinek 1989; Lu et al. 2011; Gauthier et al. 2015; Gauthier et al. 2013; Gauthier et al. 2012; Bhogal et al. 2016; De Vis et al. 2015). The mechanisms underlying this decreasing reactivity of cerebral vessels with aging are postulated to be related to local vascular stiffening (Desjardins 2015). An interesting finding of our study is that sex did not moderate the relationship between CVR and EF among older adults, suggesting that other hemodynamics mechanisms may be at play. Interestingly, recent work published by our group has shown that CVR, which could be interpreted as an indirect measure of vascular elasticity in the brain, may not be uniquely dependent on brain-based vascular properties and may be partly dependent on more global properties such as changes in endothelial function, chemosensitivity and cerebral autoregulation (Intzandt et al. 2019). Indeed, the lack of moderation between CVR and cognition could be attributable to the fact that this indirect measure of stiffness in the brain may be biased by unknown physiological changes and may underestimate the effects of arterial stiffness on cerebral hemodynamics. As such, CVR should be interpreted with caution as a surrogate measure of stiffness in the brain. Nonetheless, future work aiming to disentangle the relationships between CVR and cognition should implement more robust and direct measures of vascular elasticity in the brain (Baraghis et al. 2011; Warnert et al. 2016).

### 4.6 The Effect of Hematocrit

Additionally, we aimed to assess if the sex effects observed in the moderation effect (PWV*SEX) on EF were driven by differences in hematocrit. Our findings suggest that HCT may be in part driving these sex differences. Interestingly, it has previously been shown that males and females have different levels of hemoglobin which affect MR measures of flow (Kimberly et al. 2013;Yip et al. 1984; Vahlquist and Others 1950; Garn et al. 1975). Since PWV, a surrogate marker for stiffness, is dependent on the differences in pressure which in turn is dependent on radius of vessels and blood viscosity, it is plausible that our results partly reflect a blood viscosity effect (Stojadinović et al. 2015; Painter 2008). We reasoned that PWV as a measure of stiffness may not be pure. Interestingly, most published studies have ignored the effect of blood viscosity on PWV measurement, since the measure is based on the underlying assumption that blood viscosity is constant across subjects (Stojadinović et al. 2015; Parkhurst et al. 2012). It has been previously shown that there might be some relation between blood flow and vascular wall, and that blood viscosity as a mechanical property of blood flow might affect measures of PWV (Kim et al. 2013).

### 4.7 Limitations

The results from this study should be viewed in light of some limitations. First, menopausal status and hormonal levels were not acquired at the time of the study. It is assumed that our female population is predominantly postmenopausal given that the age range was 55-75 years, with a mean age of 63. In fact, the average age of naturally-occurring menopause in Canada is 51 years (Canadian consensus conference on menopause, 2006). Nonetheless, this assumption is speculative, and other studies should account for menopausal status to replicate and provide clarity on the mechanisms that underlie the present results. Future work should include younger subjects (40-55 years of age) to better study hormonal status in females.

While a strength of the current study is the use of non-invasive quantitative perfusion imaging to calculate cerebral blood flow and CVR, one must note that the ASL technique suffers from several limitations. The contrast afforded by the subtraction of tagged images is only a fraction of a percent of the functional MRI contrast, providing limited signal-to-noise ratio (SNR) (Golay and Petersen 2006; Badji et al. 2019). Furthermore, full coverage of the brain is typically not possible without advanced multi-band approaches, limiting the ability to draw conclusions on the entire brain (Badji et al. 2019; Golay and Petersen 2006). Finally, most ASL imaging methods are unable to determine whether the changes detected are true reflections of changes in flow or the result of alterations in transit times (Badji et al. 2019). This is especially problematic in the case of single delay ASL studies, such as the one presented here. Despite these limitations of ASL, it has consistently been shown that CBF-CVR is a more specific measure of vascular health than BOLD-CVR (Halani et al. 2015). Indeed, BOLD is more sensitive to vascular elasticity than CBF, and may be the best choice when sensitivity is desired over specificity (Halani et al. 2015). In addition to the general limitations of ASL, the post-labeling delay chosen in this study is suboptimal for older adults since it was optimized for a younger population (Badji et al. 2019;Intzandt et al. 2019). As such, with our limited SNR, it is possible that our CBF data before and during hypercapnia was underestimated. Thus, to better understand the relationship between arterial stiffness, cognition and CVR, longitudinal studies using multi-band approaches and multi-delay implementations with optimized post-labeling delays are necessary.

Another limitation to this study is our sample size. Indeed, our subset of female and male participants is small, limiting our ability to fully generalize our findings to the general population. Indeed, it may be plausible that due to our small sample size we were unable to detect the mediating effect of sex in some of our models. In addition, our ratio of males (n=17) to females (n=31) in this sample may be biased toward females, underrepresenting our male population. As such, our study may lack adequate statistical power to detect an effect size of practical importance, especially in males. Nonetheless, our analyses were bootstrapped which can overcome the power problem of small samples. Also, our sample included predominantly very healthy older adults, limiting our ability to generalize our findings to other older adults, who often suffer from cardiovascular risk factors. We speculate that associations reported here would likely be stronger in a cohort with worse cardiovascular health. Moreover, a follow-up study with better statistical power is needed to confirm our findings.

Finally, this study has a cross sectional design making it difficult to draw general conclusions on the population. Although this study provides valuable knowledge on the impact of sex on the relationship between aortic stiffness, cognition an CVR, a longitudinal study is needed to better understand the sex differences among those hemodynamic measures across the lifespan to better personalize CVD prevention strategies.

## 5. Conclusion

The findings from this study add to a growing body of research that underscores the interrelationship between cardiovascular function and brain health among aging adults. Overall, this paper identified a sex moderation between PWV and CVR, PWV and EF but not between CVR and EF in a sample of healthy older adults. Our data also demonstrated that HCT may play a role in driving some of these sex effects. These findings could be the results of different sex hormones, such as estrogen and testosterone, that are known to alter cerebrovascular measures of brain health. Finally, as the key oxygen-carrying molecule in the body, hemoglobin may play a direct role in influencing vascular measures of stiffness such as PWV and affect MR measures of flow. More importantly, understanding these hemodynamic associations may lead to earlier detection and targeted interventions to prevent or lessen the onset of cardiovascular diseases linked with higher aortic stiffening. Thus, future longitudinal studies that explore sex differences should include evaluation of the role of hemoglobin and investigate the role of hormone variations (i.e. sex hormones).

## Abbreviation

CVD: Cardiovascular disease
MRI: Magnetic resonance imaging
ASL: Arterial spin labelling
PWV: Pulse wave velocity
CVR: Cerebrovascular reactivity
HGB: Hemoglobin
HCT: Hematocrit
CSF: Cerebrospinal fluid
VBM: Voxel based morphometry
PP: Pulse pressure
AS: Arterial stiffness
CBF: Cerebral blood flow
EF: Executive function
PET: Positron-emission tomography
TCD: Transcranial doppler
PS: Processing speed
MMSE: Mini-mental state examination
CWIT: Color-word interference test
TMT-B: Trail making test part B
SNR: Signal-to-noise ratio
WMH: White matter hyperintensities
pCASL: Pseudo-continuous arterial spin labeling
FLAIR: Fluid-attenuated inversion recovery
NO: Nitric oxide

## Acknowledgements

The authors thank Carollyn Hurst and André Cyr for their help with data acquisition, Élie Mousseaux, Alban Redheuil, Muriel Lefort, Frédérique Frouin, and Alain Herment for their help with the aortic protocol and analysis, Cécile Madjar, Mélanie Renaud and Élodie Boudes for their help with logistics, and Catherine Foster for helpful discussions. The authors thank Ellen Garde, Arnold Skimminge and Pernille Iversen for their help with vascular lesion segmentation. They thank Céline Denicourt for performing the blood draws. They thank Jiongjiong Wang of the Department of Neurology at UCLA who provided the dual-echo pseudo-continuous arterial spin labeling sequence. This work was supported by the Canadian Institutes of Health Research (MOP 84378, Banting and Best Scholarship held by C.J. Gauthier), the Canada Foundation for Innovation (Leaders Opportunity Fund 17380), the Ministère du développement économique, de l’innovation et de l’exportation (PSR-SIIRI-239), the Canadian National Sciences and Engineering Research Council (R0018142, RGPIN 2015-04665), the Heart and Stroke Foundation of Canada (New Investigator Award held by C.J. Gauthier), and the Michal and Renata Hornstein Chair in Cardiovascular Imaging (Montreal Heart Institute, held by C.J.G.).

